# Evolutionary rewiring of the human regulatory network by waves of genome expansion

**DOI:** 10.1101/164434

**Authors:** Davide Marnetto, Federica Mantica, Ivan Molineris, Elena Grassi, Igor Pesando, Paolo Provero

## Abstract

Genome expansion is believed to be an important driver of the evolution of gene regulation. To investigate the role of newly arising sequence in rewiring the regulatory network we estimated the age of each region of the human genome by applying maximum parsimony to genome-wide alignments with 100 vertebrates. We then studied the age distribution of several types of functional regions, with a focus on regulatory elements. The age distribution of regulatory elements reveals the extensive use of newly formed genomic sequence in the evolution of regulatory interactions. Many transcription factors have expanded their repertoire of targets through waves of genomic expansions that can be traced to specific evolutionary times. Repeated elements contributed a major part of such expansion: many classes of such elements are enriched in binding sites of one or a few specific transcription factors, whose binding sites are localized in specific portions of the element and characterized by distinctive motif words. These features suggest that the binding sites were available as soon as the new sequence entered the genome, rather than being created later by accumulation of point mutations. By comparing the age of regulatory regions to the evolutionary shift in expression of nearby genes we show that rewiring through genome expansion played an important role in shaping the human regulatory network.

## 1 Introduction

Evolution of the regulatory network is believed to underlie a significant fraction of the phenotypic divergence between vertebrates [1–3]. Genetic events affecting gene regulation can be classified into two classes: exaptation of existing sequence through the accumulation of small-scale mutations, and de-novo appearance of regulatory DNA through genome expansion driven for example by transposable elements (TE). Both mechanisms have been shown to be relevant in the evolution of human regulatory DNA [4–10].

In particular, information-rich binding sites such as the one recognized by CTCF are much less likely to arise through the accumulation of random point mutations than simpler binding motifs: indeed it was shown [6] that the expansion of lineage specific transposable elements efficiently remodeled the CTCF regulome. The activity of TEs in generating TFBS was studied more generally in [7], where it was observed that about 20% of BS were embedded within TEs, thus revealing the latent regulatory potential of these elements [5]. The role of a specific class of TEs in generating transcription factor binding sites was recently investigated in [10]. On the other hand, it was recently shown [8] that recent enhancer evolution in mammals is largely explained by exaptation of existing, ancestral sequence rather than by the expansion of lineage-specific repeated elements. A systematic investigation of the role of genomic sequence expansion in rewiring the regulatory network is however still missing.

In previous works we investigated the most recent evolution of the human regulatory network by looking at both promoter sequence divergence [11] and genomic expansion after the split from the chimp [12]. Here we attempt to reconstruct a much longer evolutionary history, focusing on regulatory evolution through genome expansion since the common ancestor of all Vertebrates. To this aim we develop a segmentation of the human genome based on sequence age. By overlaying this segmentation onto Transcription Factor (TF) binding data we can reconstruct how successive waves of genome expansion modified the regulome of each TF.

We then examine some signatures that can help determine whether the binding sites were present at the time of the appearance of the new sequence, or were created later by progressive accumulation of point mutations. For repeated elements, these signatures include the specificity of the TFs binding each class of repeated elements, the existence of preferred locations of the binding sites within the repeated elements, and of distinctive motif words. Finally, we use comparative transcriptomics data to determine the effect of such waves on the evolution of gene expression.

## 2 Results

### 2.1 Segmentation of the human genome by sequence age

To estimate the age of each region of the human genome we used a published multiple alignment of 100 vertebrate genomes [13]. Each region was classified as present or absent in each non-human species depending on whether it aligns to sequence or to a gap in the multiple alignment: therefore each region is characterized by a present/absent binary vector of length equal to the number of non-human species in the alignment (see Methods for details).

For each region the present/absent vector was used as input to a maximum parsimony algorithm to determine the most likely (most parsimonious) history of appearance/disappearance of the region during vertebrate evolution that explains what is observed in extant species. Note that parsimony was not used to reconstruct the phylogenetic tree, which was instead fixed, but only to reconstruct the presence or absence of the sequence in each ancestral node. We thus obtained a new vector expressing the presence of the region in progressively older ancestors, from the human-chimp common ancestor to the common ancestor of all vertebrates (see Fig. 1).

**Figure 1:**
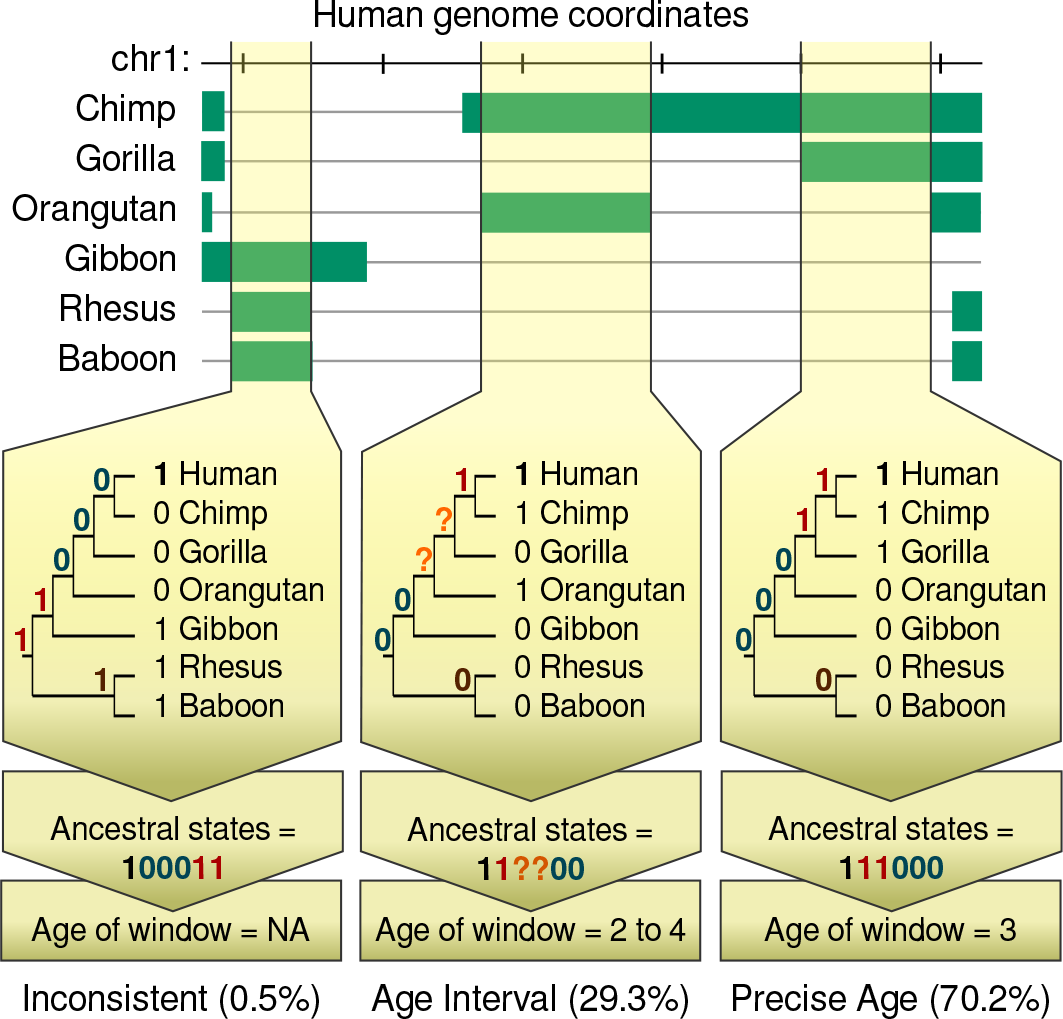
Maximum parsimony applied to a multiple alignment of the human genome to 100 vertebrates is used to infer the presence of a genomic region in each ancestor and thus the evolutionary age of the region. A green bar indicates that the human genome region aligns to sequence, rather than to a gap, in the nonhuman genome. Each region can be classified into one of three classes. “Inconsistent”: The reconstructed ancestral states are incompatible with a single birth event for the region; “Age Interval”: The reconstruction is compatible with single birth but only a time interval can be estimated; “Precise Age”: The precise time of the birth of the region can be estimated. The fraction of genome sequence falling into each class is shown.

Similar principles have been used in previous works [4, 8, 9]. All of them evaluated the age of windows of interest using the most distantly related species with an alignable sequence. The use of a parsimony algorithm allowed us to exploit in a controlled way genomic alignments with a large number of species and thus provide a segmentation of the human genome by age that is more robust and dense in terms of the human ancestors considered.

For 70% of the genome this method allowed us to determine a precise age of birth and for another 29.5% an age interval (when the parsimony algorithm reported the presence of the sequence in some ancestors as uncertain). For this latter fraction we defined as age the upper end of the interval. While this leads to a systematic overestimation of genomic ages, all results reported in the following were essentially unchanged when the opposite choice was made.

Only for 0.5% of the genome the reconstructed history was inconsistent with a single birth event, and this fraction was enriched in Low Complexity, Simple and tRNA repeats (*P <* 0.001, permutation test), possibly reflecting sequencing and alignment problems; this part of the genome was excluded from further analysis. The age segmentation of the genome thus obtained turned out to be robust with respect to the choice of the initial alignment data: using a collection of 47 pairwise “net” alignments obtained from the UCSC Genome Browser in place of the multiple alignment gave very similar results (Supp. Fig. S1).

### 2.2 The age distribution of the human genome and the role of transposable elements

The age distribution of the human genome is shown in Fig. 2A. Most of the human genome appeared after the split between placentals and marsupials: indeed only 13.5% of the human genome aligns with the opossum genome, while 43% aligns with the elephant (among the farthest eutherians from humans). The figure also shows the fraction of newly created sequence overlapping known TEs, which increases as we get closer to the present time. Older TEs are difficult to recognize today, and this probably explains at least in part their lesser prevalence in older regions of genome; nevertheless, since the ur-Boreoeutheria, the majority of newly gained sequence is still identifiable as TE. This indicates that TEs are an important driver of new sequence acquisition at least since then, and possibly further back in time. As expected, TEs are generally constant in age and their boundaries are close to age breaks, as seen in Suppl Fig. S2.

**Figure 2:**
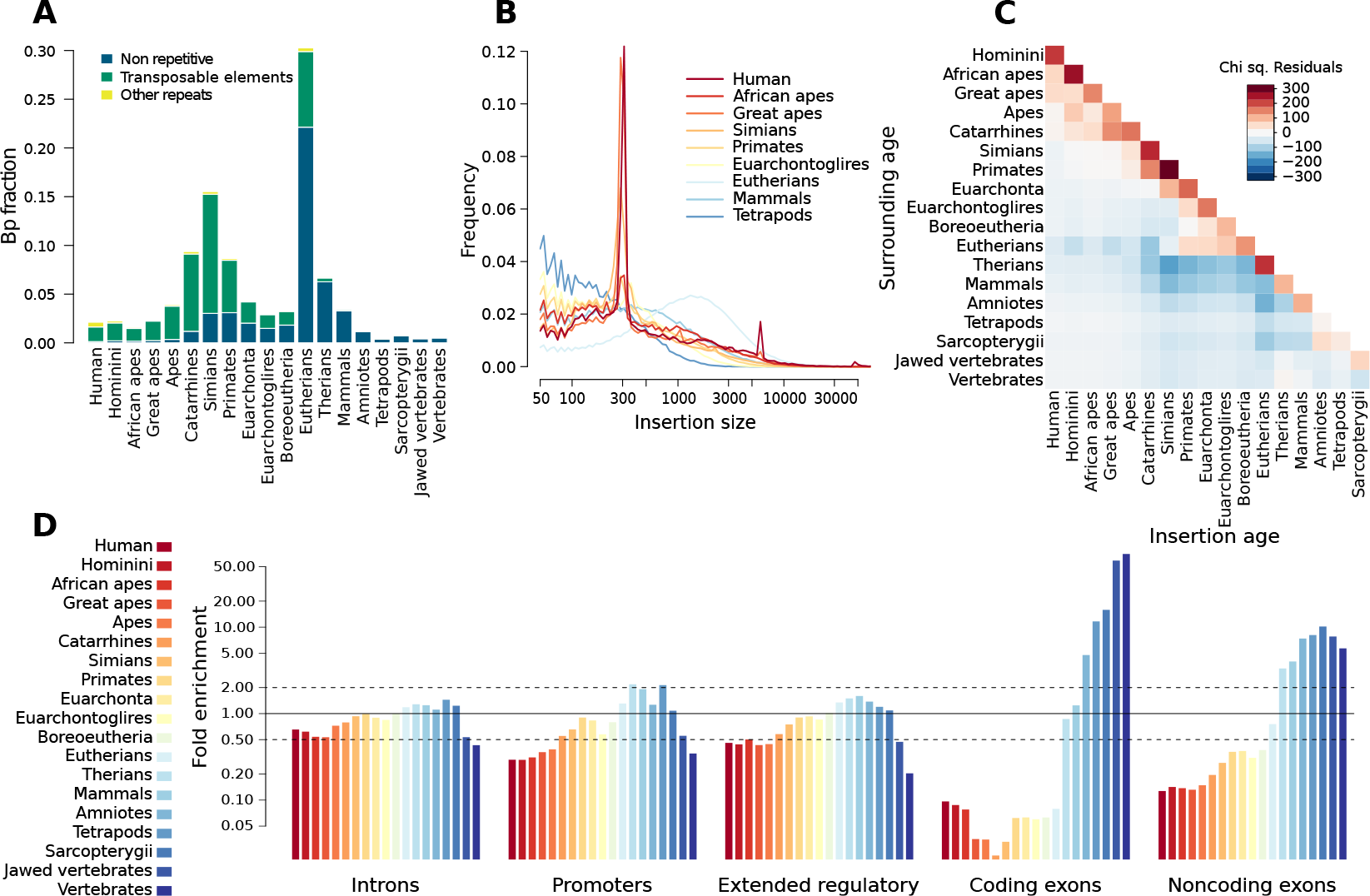
(A) Age distribution of the genome. Most of the human genome appeared after the split between placentals and marsupials. Note the increasing relative importance of TEs as we move towards younger regions. This is partly due to the difficulty in recognizing TEs that have been integrated in the genome for very long times. (B) Length distribution of reconstructed insertions of each age.Colors are associated to ages as in (A). The major peaks at length*∼*300 and*∼*6000 correspond to ALU and L1 insertions respectively (C) Comparison between the age of genomic regions (x axis) to the age of the surrounding genome (y axis) shows that new genomic sequence is preferentially inserted in younger regions, possibly because these are subjected to less selective pressure. The heatmap shows the chi squared residuals with respect to a null hypothesis of no preferential insertion, in which the insertion probability in a region of a certain age is simply proportional to the genome fraction of that age (D) Enrichment of genomic ages in functional sequence classes. We show the coverage fold enrichment compared to what expected if all functional classes shared the genome-wide age distribution.

The length distribution of reconstructed insertions of each age is shown in Fig. 2B and is driven, in particular, by waves of expansion of Alu (length*∼* 300) and L1 (*∼* 6,000) retrotransposons in the ur-Primate and ur-Homininae respectively. New insertions happen preferentially in younger regions (Fig. 2C), presumably because younger regions are subjected to weaker selective constraints, leading to the appearance of insertion hotspots within young TEs [14].

When looking at the age distribution of various functional classes of the genome we see, as expected, that age increases with functional constraint (Fig. 2D): coding exons are made by the oldest sequence, while introns are the newest. Moreover we confirmed some known results relating age to expression patterns: older genes are more expressed than younger ones [15] (see Suppl. Fig. S3A), and the coding exons of ubiquitously expressed genes are older than tissue-specific ones [16] (Suppl. Fig. S3B). However the promoters of ubiquitously expressed genes are younger than those of tissue-specific ones, perhaps due to the relaxed constraints on their fine regulation. Within tissue-specific genes the newest are expressed in testes and the oldest in the central nervous system. Importantly, the strategy by which we re-obtained these known results is completely independent from gene annotation, in contrast with the methods commonly used in classic phylostratigraphy [17, 18]. A comparison of our gene dating results with the age classes defined in [19], and those derived from the GeneTrees provided by Ensembl [20] is shown in Suppl. Fig. S4.

### 2.3 Genomic age enrichment of Transcription Factor Binding Sites

Newly acquired genomic sequence can contribute to the evolution of the regulatory network by creating binding sites for TFs [5–7]. To investigate this phenomenon in a systematic way we superimposed the results of ChIP-seq experiments performed on many TFs to the age segmentation and, for each TF, we asked whether significant age preferences could be discerned. Specifically we used a *χ*^2^ test to compare, for each TF, the number of TF binding sites (TFBS) found in each genomic age to what expected under the null hypothesis where age preference is the same for all TFs. The null model thus incorporates any deviation from the uniform distribution displayed by TFBSs as a whole, and the test reveals the specific deviations of each TF.

Out of 139 TFs, 137 showed an age distribution significantly different than the null model (*P <* 0.05 after Bonferroni correction for multiple testing), with only PPARGC1A and STAT2 not significant. TFBS local clustering could inflate the *χ*^2^ *P*-values, and can be due to technical reasons (e.g. a single binding site interpreted as multiple peaks by the peak-calling software) or biological reasons (e.g. the accumulation of multiple binding sites of the same TF in regulatory regions). These effects can be controlled by counting as a single BS peaks closer than a given cutoff. Reassuringly, the enrichment results are essentially unchanged whether we use or not a cutoff, and with cutoffs of 100 bps and 500 bps. As an alternative strategy, we replaced the *χ*^2^ *P*-values with empirical ones, by shuffling the TFBS in a block-wise manner: within each block the succession of TFBS is maintained, so as to maintain their local clustering, while blocks are randomly shuffled on the genome. This results in 135 significant TFs, excluding only two more TFs, SIRT6 and SMARCB1, compared to the *χ*^2^ analysis. These results show that most TFs show specific preferences in the age of the genomic sequence they bind. Such preferences can be visualized using the chi squared residuals, and are shown in Fig. 3 and Suppl. Figures S5 and S6. The numeric results are found in Supp. Table S1.

**Figure 3:**
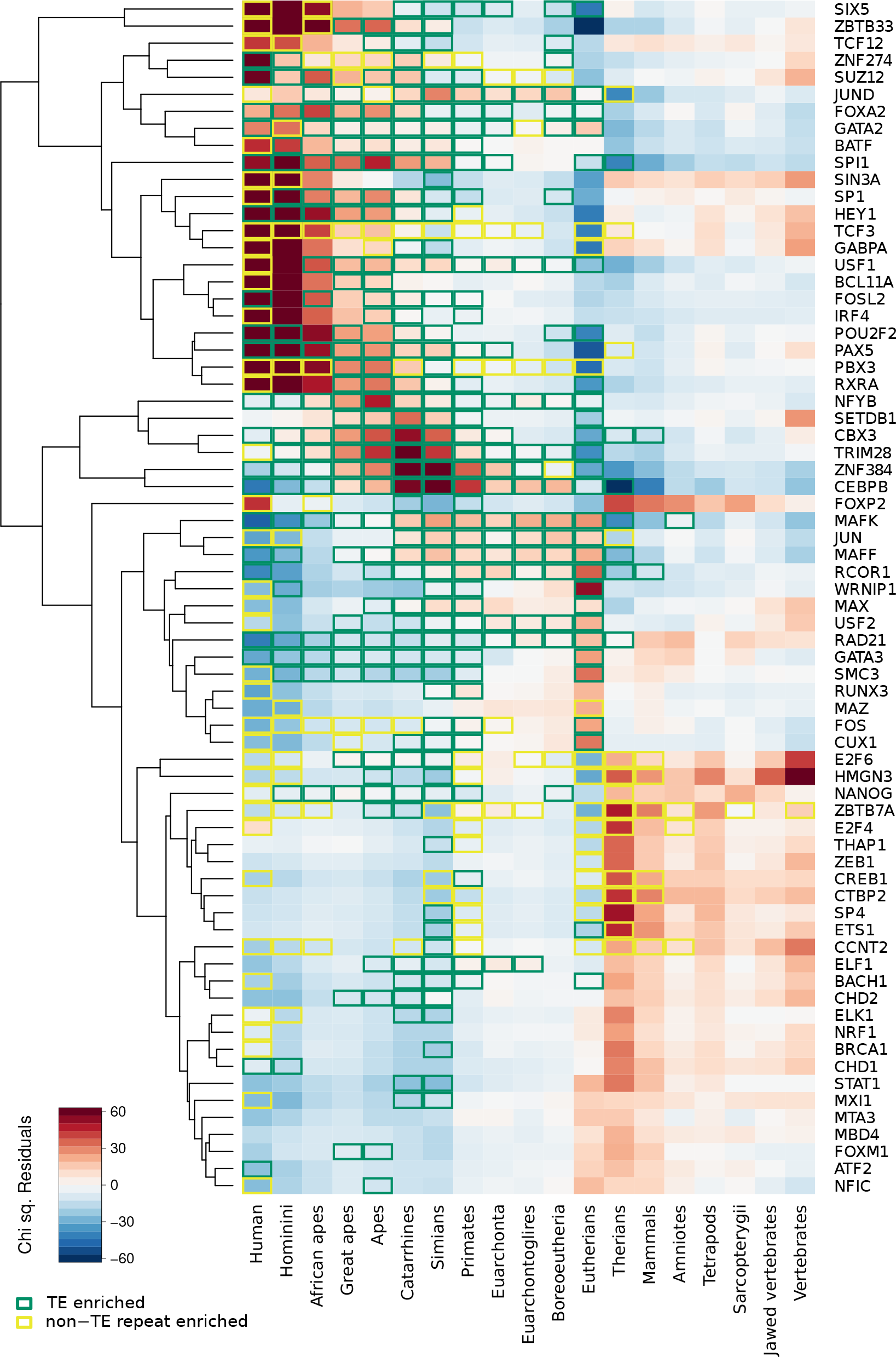
Enrichment of TFBSs in genomic regions of different evolutionary age. The heatmap represents the deviations of the age distribution of the binding sites of each TF from the overall TFBS distribution. Chi squared residuals are shown, so that positive values (red) correspond to enrichment and negative values (blue) to depletion. Only the top 50% TFs by significance are shown: a complete figure is available as Suppl. Fig. S5. TF/age combinations with significant repeat class enrichments are bordered, separately for non-transposon repeats (yellow) and transposon repeats (green).

Notably, such age enrichment in TFBS corresponds to specific functional enrichments of their target genes. In agreement with what reported by [4] for conserved non-coding regions, we observe significant TFBS enrichment near developmental and transcription factor genes in ages preceding the appearance of mammals; near receptor-binding proteins between the ur-Amniote and the ur-Eutherian; and enrichment in more recent ages near genes involved in post-translational modifications (see Suppl. Fig. S7).

### 2.4 Age enrichment suggests waves of TFBS expansions

The enrichment of binding sites of a given TF inside genomic sequence of a given age suggests an evolutionary process in which new genomic sequence extensively rewires the transcriptional regulatory network by providing existing TFs with abundant new targets. This mechanism was shown to have operated in the evolution of the CTCF regulome in mammals [6]. However, the fact that a human TFBS resides in a region that appeared at a certain time in evolution does not necessarily mean that the binding sites have the same evolutionary age. Indeed it is well known that TFBS can be generated within pre-existing sequence [8,21–23]. In this mechanism new genomic sequence could simply provide raw material for evolution to act upon by accumulation of point mutations, possibly aided by relaxed negative selection, and create TFBS that were not there when the sequence entered the genome. Other effects, such as functional characterization of genomic regions with same origin and age, could also contribute to the age enrichments shown above. In the following we will use various signatures to identify the cases in which waves of genomic expansion indeed generated an immediate rewiring of the regulatory network.

#### 2.4.1 Repeated elements carry specific motifs for specific TFs in specific portions of their sequence

If repeated elements carried TFBS at the time of their insertion in the genome, we expect to detect three signatures that are, instead, difficult to reconcile with TFBS creation by accumulation of point mutations. First, we expect each class of repetitive elements to be enriched with binding sites of just one or a few specific TFs; second, we expect such binding sites to be preferentially located in a specific portion of the repeated elements; and third, we expect the TFBS located in the RE to use a specific subset of the set of all possible motif-words (DNA *k*-mers compatible with the binding [6]).

We thus asked which of the age enrichments shown in Fig. 3 could be ascribed to the expansion of a specific, recognizable repetitive element. We considered all TF/age pairs (cells in the heatmap shown in Fig. S5) and for each of them we evaluated the number of TFBS overlapping each repeat class. We then tested whether such overlap was significantly enriched with respect to all TFBSs of the same age irrespective of the identity of the TF. That is, we asked whether the instances of a repetitive element appearing at a certain time tended to be associated to TFBSs of a specific TF.

We found a total of 3625 significant TF/RE/age triplets (Suppl. Table S2), involving 888 TF/age pairs. In Figure 3 these are shown as bordered cells. In most cases (2887 involving 600 cells) the enriched RE class is a TE. In Fig. 4 we show the TFs whose binding sites are significantly enriched in each RE class, and the distribution of the number of TFs associated to each RE class. For most classes the enrichment in binding sites are restricted to one or a few TFs.

**Figure 4:**
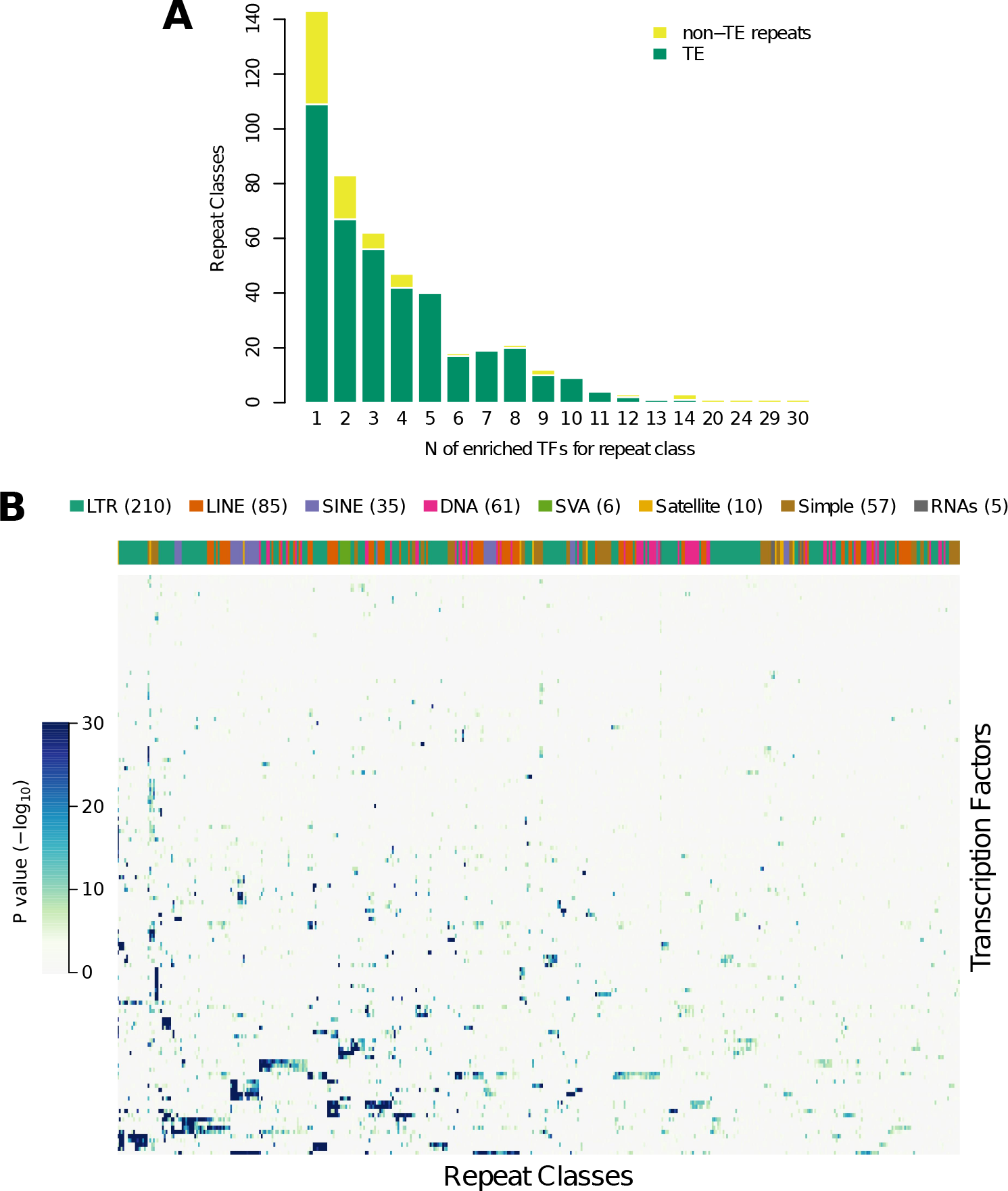
Classe of repeated elements carry specific TFs: (A) distribution across RE classes of the number of enriched TFs. (B) The heatmap shows which TFs are enriched in each class of repeated elements: for each TF-RE class pair the age with the strongest enrichment is shown, colour intensity corresponds to –*log*_10_(*P*) (Fisher’s exact test). RE classes are labeled by the top coloured bar, where each colour corresponds to a RE family. See the legend for a count of the RE classes included, divided by family.

To determine whether TFBS occur in specific positions within repeated elements we considered all the enriched TF/RE/age triplets determined above and computed the entropy of the position distribution of the TFBS (represented by the median point of the ChIP-seq peak) within instances of the RE of the appropriate age. We then used a permutation test (see Methods) to determine whether such entropy was significantly lower than that of the position distribution of all TFBS (of all TFs) on the instances of the same RE of the same age. The test thus shows whether the enriched TF is more precisely localized inside the RE compared to all TFs that bind the RE. In 1377 out of 1970 cases tested (involving 399 out of 495 TF/age combinations), we found a significantly lower entropy in the enriched TF/age/TE combination (Benjamini FDR 0.05, see Material and Methods for further details), as shown in Figure 5.

**Figure 5:**
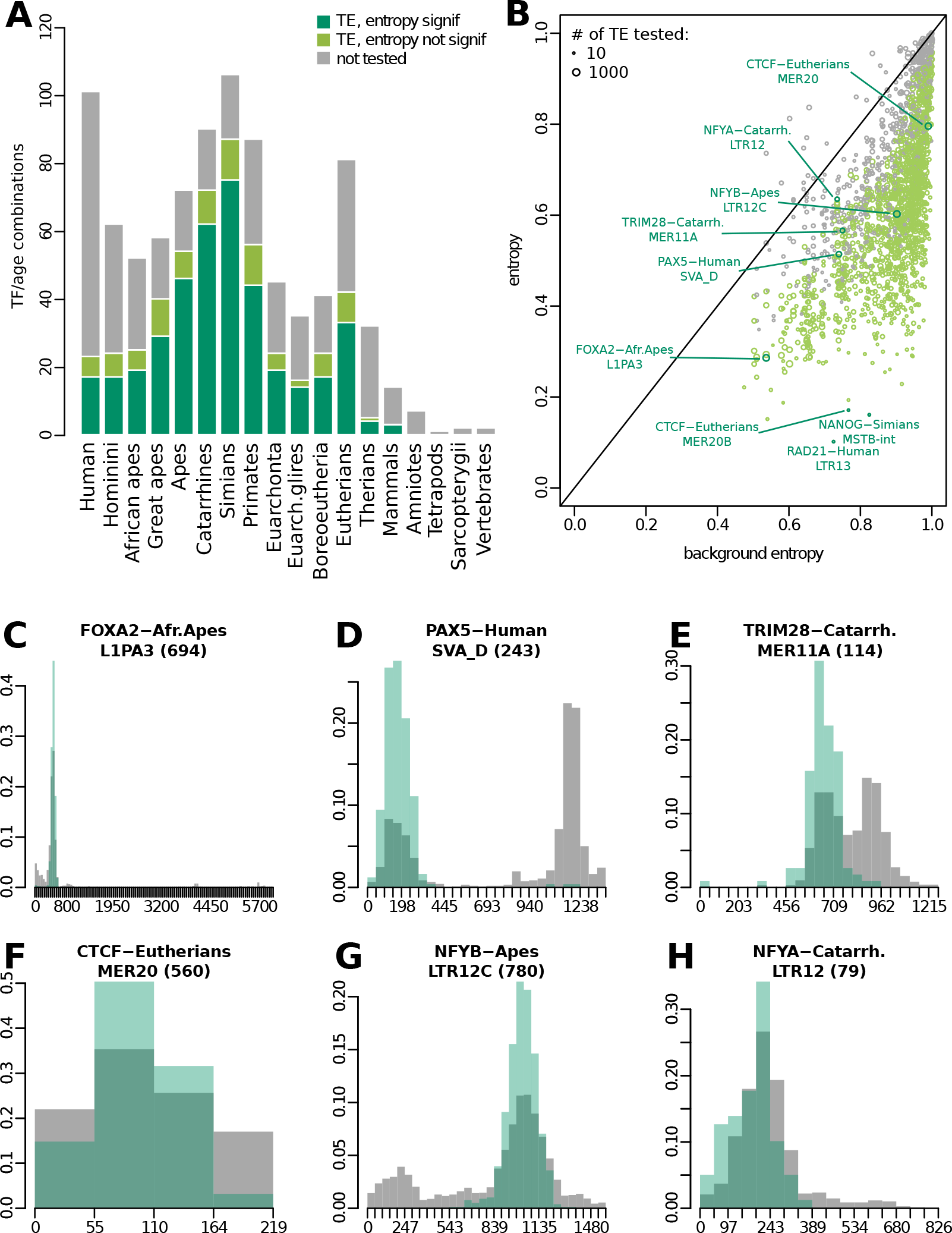
Enriched TFBS that have a constant position within Repeated elements. (A) Cases with significantly lower entropy in the position distribution with respect to the background. Each bar represents the number of TF/age pairs that resulted significant (dark green), not significant (light green), not tested (grey), for each age. (B) Entropies for each tested TF/TE/age triplet: on the vertical axis is reported the entropy of the position distribution of BS for the enriched TF within the corresponding TE, whereas on the horizontal axis the entropy of the background, that is the same distribution but considering all TF binding to the TE. Point size and colour represent respectively number of tested instances and test result, with green being significant. (C-H) Significant examples: green bars represent the position distribution of BS considering only the enriched TF, grey bars represent the background. In particular (G) is the case with most instances and (H) is being considered a borderline case with FDR 0.05.

Considering again the set of enriched TF/RE/age triplets, we then examined if binding sites lying on the enriched RE used a specific set of motif-words with respect to binding motifs placed elsewhere, similarly to what is shown for CTCF in Ref. [6]. To do so we identified the top scoring motif within all ChIP-seq peaks with a Positional Weight Matrix corresponding to the TF of interest and annotated whether the peak was falling on the enriched RE class or not. We then asked wether the two sets of TFBS differed in word composition: 523 out of 1707 TF/RE/age triplets tested (199 out of 366 TF/age combinations) were found significant against 100,000 simulations (Benjamini FDR 0.05, see Material and Methods for further details), as shown in Figure 6A.

**Figure 6:**
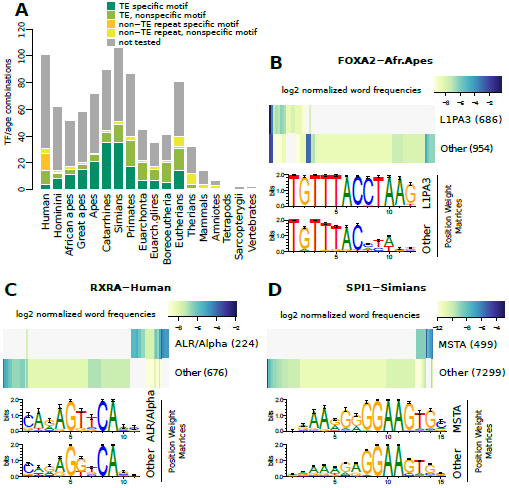
TFBS with RE-specific motif-word distribution. (A) Cases with word occurrences in the enriched RE class significantly different than elsewhere. Each bar represents for each age the number of TF/RE pairs that resulted significant (dark colors), not significant (light colors), not tested (grey), separating for transposon (green) and non-transposon (yellow) repeats. (B-D) Significant examples: the word frequency matrix is represented by the heatmap (colors are in log2 scale, white means 0 occurrences); the Positional Weight Matrix logo obtained from binding site motifs occurring on the tested RE class is compared with the logo from all other binding sites.

Altogether, in most cases of RE enriched in the binding sites of a TF we observe that the enrichment is not only specific for the identity of the TF, but also for the position of the binding within the RE and/or the word composition of the binding motif. Such specificity is difficult to reconcile with a process in which point mutations create binding sites in the ages after the appearance of the new sequence, and suggests instead that at least the genetic component of the regulatory rewiring was effected directly by the insertion of new sequence.

#### 2.4.2 Network rewiring by genome expansion causes gene expression evolution

An important question is whether the TFBSs created by waves of genomic expansion are functional, that is whether they lead to changes in gene expression. The authors of Ref. [24] used comparative RNA-seq data to identify genes whose expression underwent a shift at a certain point of the evolutionary history of mammals. By comparing the timing of expression shifts to the age of surrounding TFBSs we can investigate the role of genomic expansion in effecting gene expression evolution.

We considered all genes which, according to [24], underwent an expression shift in exactly one human ancestor (except ur-Mammal and ur-Hominoidea for which the shift cannot be attributed to a branch in Ref. [24]). We determined their associated regulatory region using GREAT [25] and, in such region, counted the TFBSs falling within each genomic age (Fig. 7).

**Figure 7:**
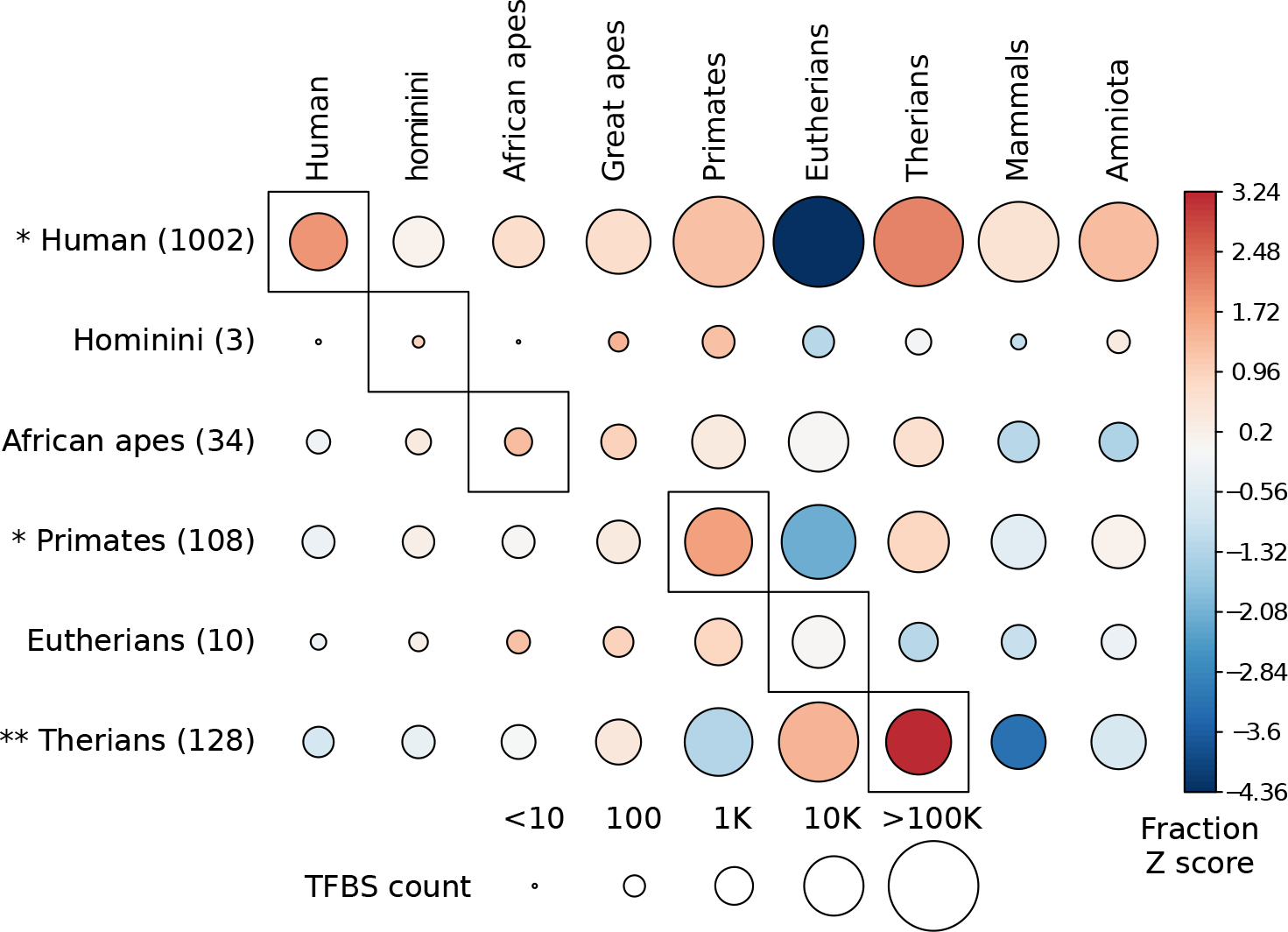
(A)Age distribution of TFBSs in the regulatory region of genes which underwent an expression shift in a human ancestor. The heatmap represents, for each ancestor in which the shift occurred, the fraction of TFBS with a certain age transformed into *z*-scores. The size of the circle represents the number of TFBS in each cell. Next to each ancestor name is annotated in parentheses the number of genes whose expression shifted. Asterisks indicate significant enrichment of TFBSs of the same age as the expression shift (*: P ¡0.05; **: P ¡0.01). *z*-scores and enrichment *P*-values were based on 5,000 randomizations of the age of the expression shift.

Such counts were then compared to 5,000 randomizations of the age of the gene expression shift, and thus transformed into *z* scores, shown in Fig. 7. Given a genomic age *a_G_* and an expression shift age *a_E_*, *z*(*a_G_, a_E_*) is positive (negative) when TFBSs of age *a_G_* are enriched (depleted) in the regulatory region of genes which shifted their expression in *a_E_*. For the three evolutionary ages with the largest number of expression shifts (Human, Primates and Therians) we observed a statistically significant enrichment of TFBS with *a_E_* = *a_G_* (permutation test based on the randomizations described above). These results show that the rewiring of regulatory networks by newly acquired genomic sequence results in an immediate, detectable change in the transcriptome.

### 2.5 Transcription factors that underwent binding site expansion

By looking at Fig. 3 we can observe several examples of transcription factors which gained targets through RE expansion. A sizable cluster of TFs preferentially bind Hominini-specific regions (that is, originating in the human-chimp common ancestor). These include immune-related TFs such as IRF4 [26], PAX5 [27] and POU2F2 [28] and TFs involved in the regulation of metabolism, such as RXRA [29, 30] (see Fig. 6C) and USF1 [31]. A few of these TFs gained new targets through TE expansion, including PAX5 (see Fig. 5D), FOXA2(see Figures 5C and 6B), SPI1 (see Fig. 6D), HEY1, GATA2; see also Fig. 5C,D.

The expansion of TRIM28 (aka KAP1) binding sites during primate evolution is also evident: its newly formed binding sites in the ur-Simiformes, ur-Catarrhine and ur-Hominoidea are preferentially found within TEs of various classes (especially LTR) in agreement with its role in repressing newly arising TEs [32], see also 5E. Among the TFs showing a marked target expansion in the ur-Eutherian we find CTCF and MAFF, which is involved in parturition by regulating the oxytocin receptor [33]. In agreement with Ref. [6], CTCF shows both ancestral binding sites predating the origin of vertebrates and several waves of expansion in mammals. Our results confirm in particular a role of MER20 and MER20B in the expansion of CTCF sites in the ur-Eutherian, as previously reported [34], see Fig. 5F.

FOXP2, a TF involved in speech and language which underwent significant mutations in the human lineage after the split with the chimp [35], shows a strong enrichment of TFBS in human-specific regions, suggesting that the evolution of its coding region was accompanied by the acquisition of new targets through binding to newly acquired genomic sequence. These new binding sites appear to occur more than expected on satellite repeats ALR/Alpha and HSATII.

## 3 Discussion and conclusions

We classified the human genomic sequence based on its evolutionary age, that is the ancestor in which the sequence first appeared. We then examined the age of regulatory sequences, defined as transcription factor binding sites determined by ChIP-seq experiments. Many transcription factors appear to have acquired new binding sites through genomic expansion, a fact that was known for some of them [6] but that we could establish in a systematic way.

Transposable elements play a crucial role in generating these waves of regulatory expansions, and in many cases specific families of them can be associated to specific waves of expansion, especially when these are relatively recent so that the originating TE can still be recognized. However our approach does not rely on databases of TE sequence, and thus is able to identify ancient waves of expansion such as the ones involving several transcription factors in the ur-Therian.

Several features of the TFBS located in repeated elements suggest they were already present at the time of genomic insertion: binding sites of specific TFs appear in fixed positions inside each class of repeated elements and use a distinctive set of motif-words, a pattern difficult to reconcile with a process in which binding sites are formed by the gradual accumulation of point mutations.

While our results suggest that most transcription factors obtained a relevant part of their binding sites during specific waves of genomic expansion, they do not imply that *most* binding sites are generated in this way. For example, only about 8.4% of all the TFBSs used to build Fig. 3 contribute to the enrichment of a TF/RE/age triplet, and can thus be specifically attributed to the expansion of the RE in a specific evolutionary age. This must be considered as a lower bound since our strict control of false positives in evaluating enrichment certainly leads to many false negatives. However, this relatively low percentage shows that our results are not in contradiction with those of Ref. [8] where it is shown that most regulatory elements are created by exaptation of existing sequence: while this is the dominant mechanism, most human transcription factors have also undergone waves of rapid target expansion driven by newly acquired genomic sequence.

Between the exaptation of pre-existing sequence and the immediate recruitment of new sequence for regulatory rewiring, various intermediate scenarios are possible, including for example new sequence carrying quasi-binding sites needing some point mutations to become effective, or binding sites that are not immediately effective because they reside within closed chromatin. Our age-enrichment map (Fig. 3) undoubtedly reflects also some of these intermediate cases.

## 4 Methods

### 4.1 Classifying the human genome based on sequence age

For each base of the human genome (hg19 release) we assigned presence/absence in modern vertebrates using the Multiz 100-way aligment [13]. A base of the human genome is present in another species if it aligns to sequence (regardless of matching) rather than a gap in such species. We defined regions of the genome as maximal stretches of consecutive bases sharing the same presence/absence vector. The state of each region in ancestral nodes was reconstructed through a parsimony algorithm on a fixed tree. The tree was obtained from the Multiz 100-way documentation, and led to the definition of 19 ancestral nodes ranging from *H. sapiens* to the ur-Vertebrate.

For each region, the parsimony algorithm uses as input the present/absent state of the region in each of the extant aligned species and returns a present/absent/unknown state in each ancestral node. We defined the age of the region as the oldest ancestor where the algorithm returned a non-absent state (present or unknown). All analyses were repeated with the opposite choice (age as the oldest ancestor with a present state) to check that our results do not depend on such bias.

For most of the genome regions (99.5% of the sequence) the reconstructed history is consistent with a single birth event (that is the region is reconstructed as absent in all ancestors older than its assigned age). The regions for which this did not happen were discarded. We also removed from our genome all regions annotated within the Gap track (downloaded on 11/22/13) in the UCSC database.

### 4.2 Overlap with repeated sequence and preferential insertions

For Repeat Elements annotation we used the Repeat Masker (downloaded on 11/19/12) and Simple Repeat (downloaded on 10/20/15) tracks, from UCSC database. We labeled as ‘Transposable Elements’ those belonging to DNA, LINE, LTR, Other, and SINE classes. The overlap of inconsistently reconstructed regions to repeat classes was tested against 1000 randomizations obtained by shuffling their genome positions.

To test whether insertions happen preferentially into recent genome we removed all regions smaller than 50bp and we defined as insertions all regions such that the two flanking regions are older and of equal age. We then counted the insertions happening inside regions of all possible ages, and tested their distribution against the null model in which the probability of insertion into a given age is proportional to the total genomic sequence of that age, using a *χ*^2^ test.

### 4.3 Evolutionary age of functional classes of sequence

To each RefSeq gene we attributed five functional classes of sequence: coding exons, non-coding exons and introns were defined based on RefSeq annotation. To define regulatory classes we used the genome segmentation of Ref. [36]: we consider as regulatory sequence all sequence that is not inside an exon and falls into one of the regulatory classes of [36] (classes 1-8) in at least one cell line, after removing the cancer derived cell lines Hepg2 and K562. We then classified as *promoters* all regulatory sequence within -5kbp and +1kbp of a RefSeq TSS and *extended regulatory* all regulatory sequence within the extended regulatory region defined associated by GREAT [25] to the gene.

### 4.4 Evolutionary age and gene expression

We used the expression profiling of human tissues from the RNA-seq Atlas [37] and we defined a gene as expressed in a tissue if its RPKM was *>* 1. We then compared the mean genomic age of tissue-specific or ubiquitary genes vs. all other genes with a Wilcoxon-Mann-Withney test, separately for each sequence functional class. We also evaluated the Spearman correlation between mean genomic age and gene expression expressed as RPKM.

### 4.5 TFBS distribution in age

To investigate the age distribution of TFBS we collected all ENCODE ChIP-seq datasets, merged all the binding sites of the same TF from different cell lines and removed data from non-sequence-specific binding events (PolII and general transcription machinery) or non endogenous (GFP-conjugated proteins).

We removed peaks wider than 5kb and set a minimum peak width of 100bp, enlarging narrower peaks up to this size. We repeated the analysis with 0 or 500 minimum peak width cutoff. To each TFBS we attributed the age of its median point, after discarding all peaks overlapping a narrower peak. We then compared the age distribution of the binding sites of each TF to a null model defined by the age distribution of all TFBSs taken together, using a *χ*^2^ test. The chi squared residuals are visualized in Fig. 3 and Suppl. Fig. S5 and represent the enrichment of binding sites of a given TF in regions of each evolutionary age. To generate an empirical *P*-value, we performed 5000 times the following randomization: divide the genome into (unequal) windows, each containing exactly 200 TFBS; each window defines a sequence of 200 TF names; randomly permute these sequences of names among the windows. In this way the TFs are assigned randomly to genomic regions while conserving the local clustering of binding sites of the same TF.

### 4.6 TFBS enrichment in GO categories

TFBS were associated to genes using GREAT [25]. We evaluated the enrichment of TFBS of each evolutionary age in genes belonging to GOs where specific peaks of regulatory innovations had been identified in Ref. [4]: GO:0003700 (transcription factor activity), GO:0032502 (developmental process), GO:0005102 (receptor binding) and GO:0043687 (post-translational protein modification). The enrichment was defined as the fold enrichment of the number of TFBS of each age associated to genes in each GO category compared to the number of TFBSs of the same age regardless of GO association.

### 4.7 TFBS enrichment in RE classes

To evaluate the enrichment of RE classes we used Fisher’s exact tests: for each TF/age/RE combination, we tested whether the binding sites of the specific TF tend to overlap instances of the RE more often than binding sites of the same age irrespective of the identitiy of the bound TF. *P*-values were Bonferroni corrected. Repeat coordinates were obtained from RepeatMasker.

### 4.8 TFBS position in REs

We kept all REs overlapping a TFBS, of length between 80% and 120% of the RE model length obtained from RepeatMasker metadata, discarding shorter and longer instances. We tested only those TF/age/RE combinations which after this filter retained at least 10 instances: only transposon classes survived after this process. We annotated the positions of the TFBS peak median point, normalized them over the length of the TE instance, and built a distribution with bins of about 50bp: the precise size of the bin ensured that the TE model could be divided into bins of equal length. For each TF/age/TE combination, call *A* the set of the binding sites of any TF falling in the specified age and overlapping the RE, and *S* its subset where the TF is the one under investigation. We computed the entropy of the binned position distribution of *S* and compared it to 10,000 random subsamplings of *A*, each of size equal to #*S*.

### 4.9 TFBS word composition

Using the JASPAR Core Vertebrates set [38], we were able to associate a Positional Weight Matrix to 66 of the 127 TFs to be tested. For each TF/age/RE triplet to be tested, we searched each Chip-seq peak with the corresponding PWM, keeping the top scoring site and discarding peaks where such site scored less than half of the highest possible score. Accounting separately for TFBS falling on instances of the enriched RE class and for TFBS placed elsewhere, we computed a contingency matrix for all occurring words and obtained a P-value for it with a Monte Carlo simulation with 100,000 replicates, using *χ*^2^ as the test statistic.

### 4.10 TFBS and gene expression shifts

TFBSs determined as above were associated to genes using GREAT [25]. For all the genes which underwent an expression shift in exactly one human ancestor (in any tissue) according to Ref. [24] we counted the associated TFBSs of each evolutionary age. These counts were compared to those obtained in the same way after randomly shuffling 5000 times the age of the expression shift and thus transformed into *z*-scores shown in Fig. 7. Statistical significance of the enrichment of the diagonal element of each row was determined empirically by comparing the number of TFBSs of the same age as the expression shift to the distribution of the same number in the 5,000 randomizations.

## 5 Authors contributions

Conceived and planned the study: DM, PP. Performed computational analsyses: DM, FM, IM, EG, IP, PP. Wrote the manuscript: DM, PP. All authors read and approved the final manuscript.

## 6 Acknowledgements

We would like to acknowledge insightful discussions with Ugo Ala, Davide Cittaro, Tyler Linderoth and Emilia Huerta-S`anchez.

---

**Figure S1:**
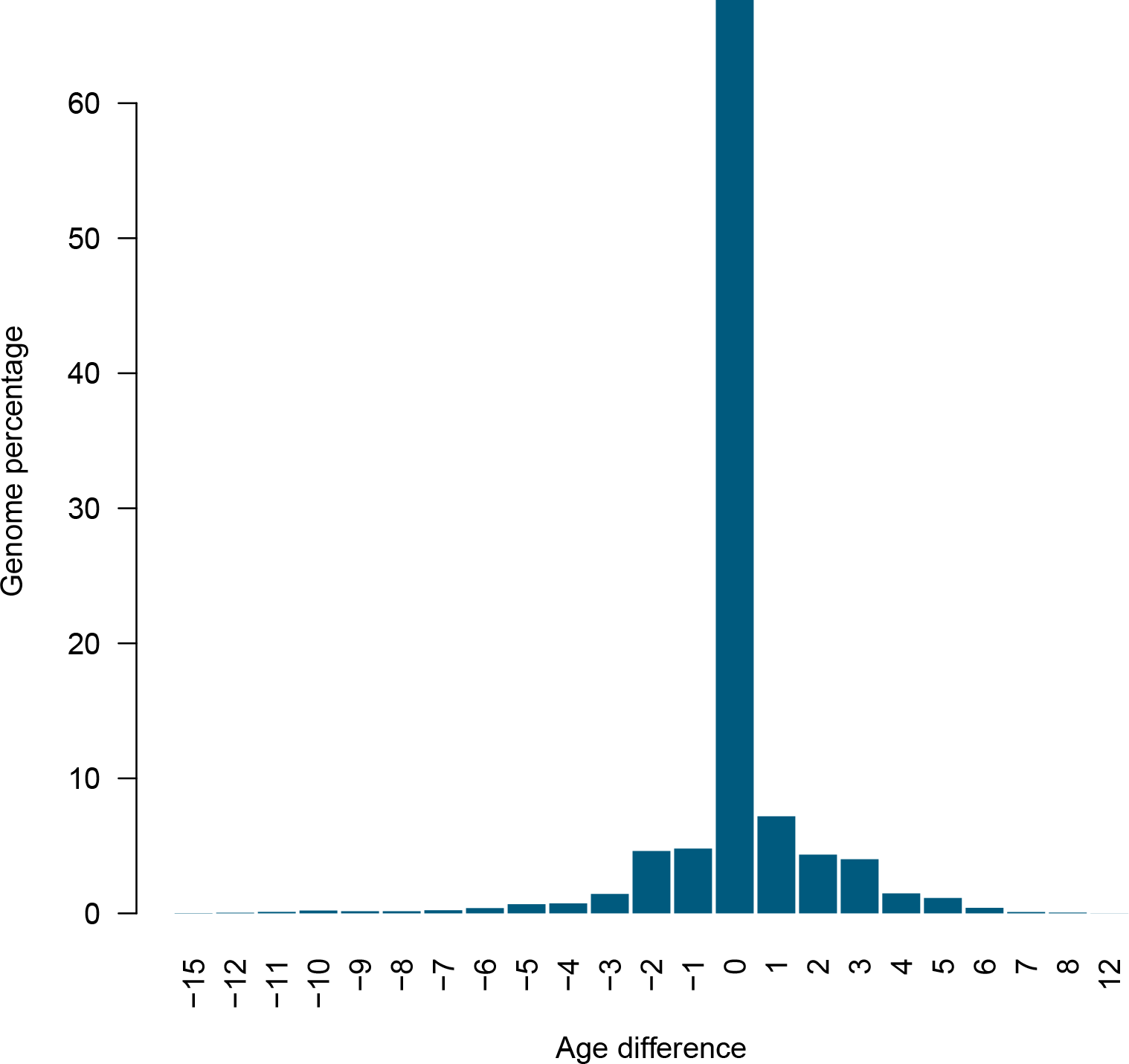
Distribution of the difference between genomic ages evaluated starting with multiple alignments or pairwise alignments between the human genome and 100 vertebrate genomes.

**Figure S2:**
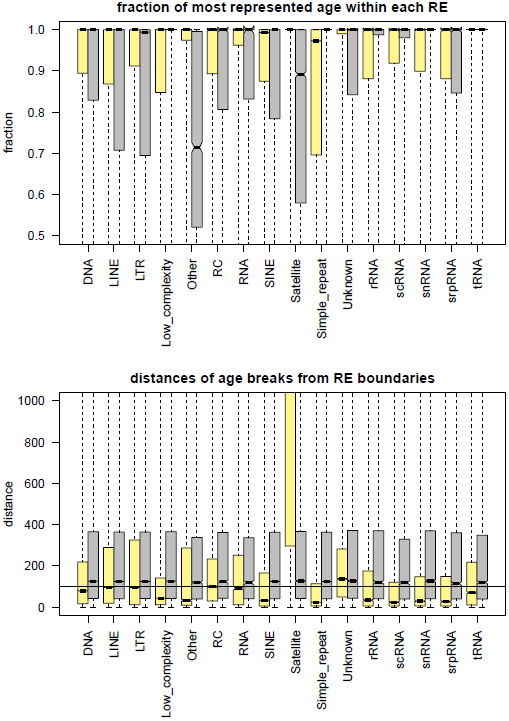
Identity of Repetitive Elements and age blocks. Repeats are grouped for class. Each repeat class (yellow) is compared to a random set of regions with same size and length distribution (grey). (A) Distributions of fraction covered by the most represented age within each element: RE tend to have constant age. (B) Distributions of distances between repeats boundaries and age breaks. Except for Satellites and Unknown repeats, age breaks and RE borders tend to appear closer than 100 bp and with respect to their randomized counterpart.

**Figure S3:**
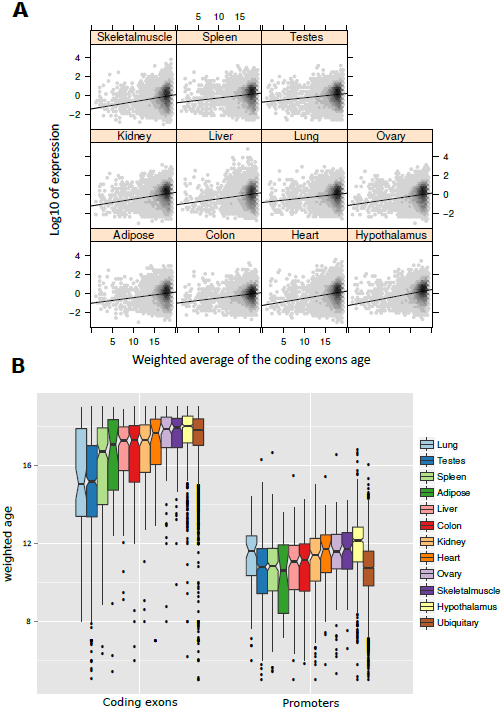
Gene expression and gene age. The age of a sequence is defined as the average age of the regions overlapping the sequence, weigthed by overlap length. (A) Older genes are more expressed: the panels show the dependence of gene expression in a collection of human tissues [37] from the age of their exonic sequence. (B) Ubiquitously expressed genes are older in their coding exons but younger in their promoters than tissue-specific genes.

**Figure S4:**
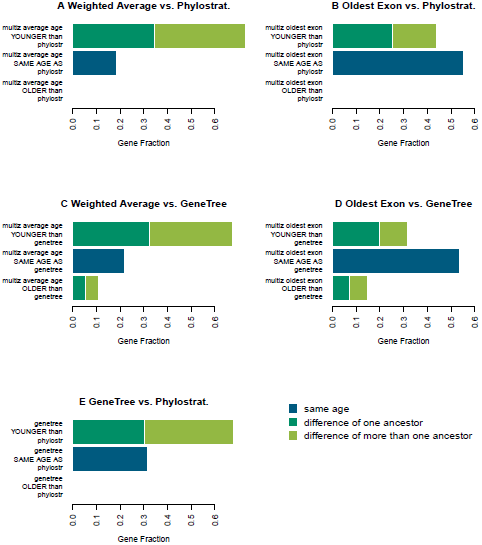
Comparisons between gene age assigned by different dating methods. Weighted Average is defined as the average age of the regions overlapping the gene, weigthed by overlap length, while Oldest Exon is defined as the age of the oldest sequence at least 100 bp long within the gene. Only coding regions are considered. **Phylostrat**: Age is obtained from [19] by translating older age classes into our oldest age (Vertebrates). **GeneTree**: age is obtained from Ensemble GeneTrees, dating each gene to the last common ancestor of human and the most distant species in which any type of ortholog was detectable. When we used the Oldest Exon approach the majority of genes showed the same age attributed by Phylostrat or GeneTree, reflecting the ancestor in which the core of the gene was generated de novo. On the other hand the Weighted Average tend to assign more recent ages: this age definition decreases when either new sequences are acquired or existing exons are lost.

**Figure S5:**
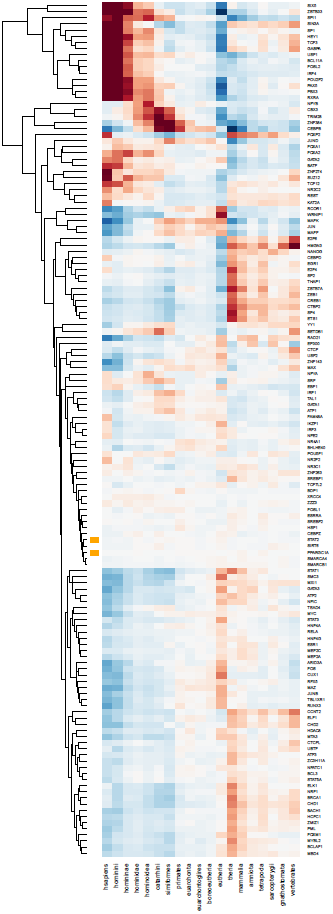
Same as Fig. 3, including all TFs studied

**Figure S6:**
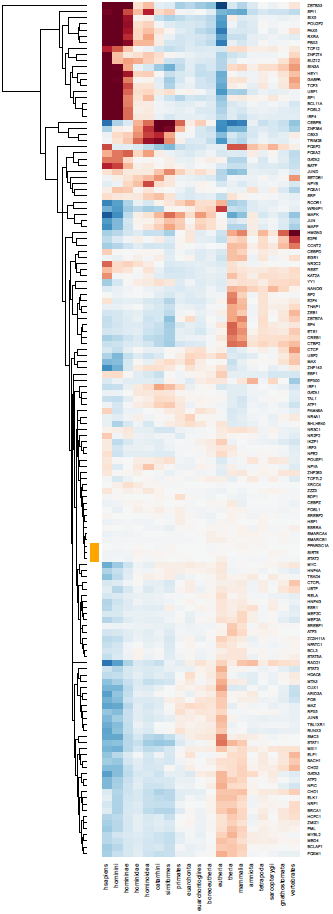
Same as Supp. Fig. S5, but based on minimum genomic ages

**Figure S7:**
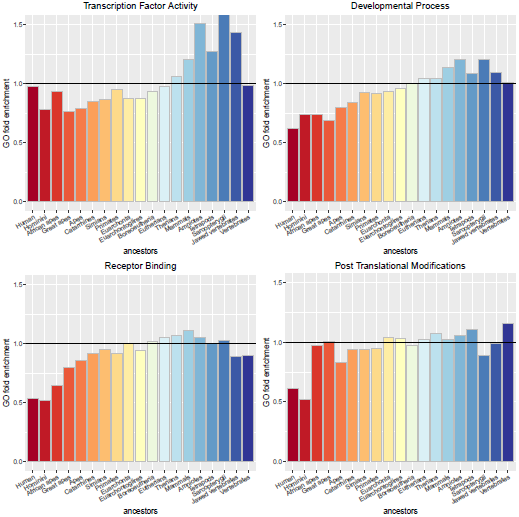
Age enrichment of TFBSs targeting genes in the GO categories shown in Ref. [4]. We show the ratio between the number of TFBS of each age targeting genes annotated to the GO category and the expected number based on the total number of TFBS of the same age.

**Figure S8:**
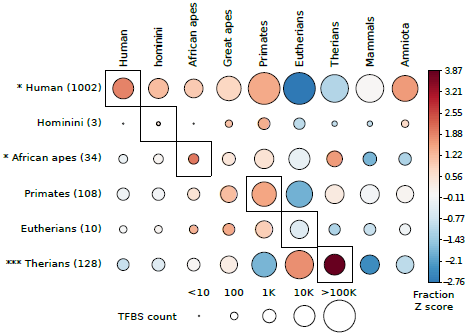
Same as Fig. 7, but based on minimum genomic ages

Table S1: Chi squared *P*-values and residuals for all TF/age combinations, used to build the heatmap in Figs. 3 and S5

Table S2: Repeat element classes significantly associated to transcription factor binding sites of a specific age. We report the TE/TF pairs whose overlap is significantly enriched with respect to all TFBS of the same age. The columns represent: Transcription Factor (TF); Repeated Element (RE); Age of enrichment (age); Phylum whose common ancestor corresponds to the age (Phylum); Fold enrichment and P-value with respect to TFBS of the same age (FC and P)

